# DEVELOPMENT AND EVALUATION OF COATED RICOTTA CHEESE WITH WHEY PROTEIN AND CLOVE OIL

**DOI:** 10.1101/2023.10.25.563938

**Authors:** Muhammad Aamir, Eram Sarwar, Aftab Ahmad, Farhan Saeed, Ali Ikram, Muhammad Afzaal, Faiza Kiran, Huda Ateeq, Noor Akram, Shahzad Hussain, Mahbubur Rehman Khan

**Author notes:** **Corresponding author 1 Mahbubur Rehman Khan**, Department of Food Processing and Preservation, Hajee Mohammad Danesh Science & Technology University, Dinajpur-5200, Bangladesh. **Corresponding author 2 Muhammad Afzaal**, Department of Food Science, Government College University Faisalabad, Pakistan.

## Abstract

Cheese, oldest dairy product which is used to preserve the nutrients of milk for long time. It is available in different shapes, sizes, flavors and textures. Ricotta is fresh soft cheese, is prepared through coagulating the whey proteins. It is a value-added product manufactured from cheese whey. Microbial spoiling occurs as a result of unmanaged conditions, resulting in the unfavorable changes in odor, flavor, and texture. That leads to food poisoning when infected food is consumed. Natural preservatives are preferred by consumers over synthetic preservatives since they are safer and extend the shelf life of food goods. Synthetic preservatives are poisonous and have negative health consequences. Edible coatings are biodegradable, natural films that can improve the product’s safety, quality, and nutritional content until it is consumed. This research was carried out to analyse the effect of whey protein and clove essential oil on the shelf life of ricotta cheese due to their excellent gas barrier properties, antioxidant and antimicrobial properties. Different concentrations of clove oil (0.1, 0.2, 0.3 %) are incorporated in whey protein coating solution (2.5,5,7.5%). The overall findings for all groups showed that there was gradual decrease in acidity and moisture content. Therefore, sample T_RC3_ treated with 0.75 % whey protein and 0.3% clove oil showed significant results (P<0.05). They showed less microbial count, increased fat and protein value. But it also had great impact on sensory analysis such as on color, texture, aroma while it showed significant overall acceptability due to high clove oil and whey protein concentrations. Hence, this study showed significant effect on shelf life of ricotta cheese and its shelf life increased from 7 to 21 days.

## 1. INTRODUCTION

By definition, milk is a nutrient-rich liquid that makes a good culture medium for a variety of species. It provides a variety of nutrients, including vitamins, proteins, lipids, and carbs, supporting a wide range of microorganisms in ideal nutritional conditions (Rollins et al., 2023). Milk is known as a complete food because of its high nutritional value. Milk is consumed by about 6 billion people all over the world (Thorning *et al.,* 2016). It is used to make yogurt, cheese, butter, and other dairy products which are widely consumed due to their high calcium levels (Boutron-Ruault and Bonnet 2020).

Cheese is a concentrated form of proteins and lipids. It also enhances shelf life of nutrients. It is made from milk by coagulating the protein, removing the whey, and allowing the curd to develop naturally or using any coagulant under controlled conditions. It is one of the most nutrient rich dairy products in the world, both in terms of quantity and diversity. Almost 1000 distinct types of cheese are produced around the world (Youssef *et al.,* 2017). Cheese is a nutrient-dense food which is more susceptible to deterioration due to its biological, physical, and chemical factors. Ricotta is fresh soft cheese, prepared through coagulating the whey proteins. It’s a dairy product that is ready to eat and has a subtle sweet taste. Due to its excellent nutritional content and medicinal function, its intake is increasing every day (Hamdy *et al.,* 2018). It is healthy because it has more protein content and a creamier texture than ricotta cheese with cheese whey. Fresh ricotta cheese has a high moisture level and pH is about 6.0. It has a water activity of 0.974-0.991 and low salt concentrations, making it susceptible to microbial deterioration. As a result, even at 4 ^0^C, it has a short shelf life (Fernández *et al.,* 2014). Its pH ranges from 6.10 to 6.80. Even when kept in the refrigerator, the shelf life of fresh ricotta, a dairy cheese, is just two to three days (Ricciardi et al., 2022)

Various factors diminish its shelf life, including enzymatic degradation, weight loss, lipid oxidation, and mesophilic and psychotropic bacterial growth. Preservatives, modified atmospheric packing (MAP), high pressure, active coatings, natural and edible coatings, and a combination of technologies can all be used to extend cheese shelf life (Repajic *et al.,* 2019).

Microbial spoiling occurs as a result of unmanaged conditions, resulting in unfavorable changes in odor, flavor, and texture. That leads to food poisoning when infected food is consumed. According to reports, L. monocytogenes can thrive in whey cheeses, and even initially modest levels of contamination can result in concentrations that could be dangerous to human health. There is proof that L. monocytogenes may contamination whey cheeses and that certain cheeses encourage its growth, even when kept refrigerated (Bintsis & Papademas, 2023). Natural preservatives are preferred by consumers over synthetic preservatives since they are safer and extend the shelf life of food goods (Arshad *et al.,* 2020).

Before ripening and during storage, cheese quality defects can be controlled by applying a coating to the surface of the cheese. It also extends cheese’s shelf life. In the cheese business, consumer preference plays an important role in packaging and product coating selection. Food goods are traditionally wrapped in non-biodegradable packaging materials such as glass, metals, and plastics, which pollute the environment (Youssef *et al.,* 2017). Edible coatings are biodegradable, natural films that can improve the product’s safety, quality, and nutritional content until it is consumed. Choosing the right cheese covering is also a huge difficulty for the cheese sector in order to control quality defects When combined with other biopolymers such as carbohydrates and proteins to form a biopolymer such as polysaccharides (carbohydrates and gums as a bonus, they improve the cheese’s texture and protect the cheese’s skin from drying out, microbial contamination and chemical alterations. In order to extend the shelf life of cheese, several coatings are utilized, including chitosan, whey protein, alginates, carrageenan, galactomannans, carboxymethylcellulose (CMC), starch, and starch derivatives, to name a few (Youssef *et al*., 2017). Many fresh cheeses are coated with antimicrobial edible coatings to inhibit post processing contamination. The present study aimed to evaluate the physicochemical, nutritional and antioxidant properties of ricotta cheese coated with whey protein and clove oil and to check the effect of whey and clove oil on the shelf life of ricotta cheese.

## 2. MATERIALS AND METHODS

The research on ricotta cheese development and its coating effect on shelf life was performed in Dairy Laboratory of National Institute of Food Science and Technology, University of Agriculture, Fsd. For preparation of ricotta cheese fresh milk was procured from dairy farm, University of Agriculture, Fsd. Clove oil, and whey protein powder were bought in Faisalabad’s local market. Cloves were used as an active ingredient in the coating solution, which was then kept refrigerated (4°C).

### 2.1. Physicochemical analysis of milk and whey

The pH of milk and whey was checked by the method described by AOAC (2016). For this purpose, a pH meter (Hanna HI 99163) was taken. The titratable acidity of milk and whey was analyzed through titration method as described by Suliman *et al*. (2019). Total soluble contents of the ricotta cheese were determined by using refractometer according to the method as explained in AOAC 2016. Milk and whey sample’s moisture content was analyzed by using moisture analyzer by following the procedure as explained by Ríos-de-Benito et al. (2021). Protein contents of fresh milk and cheese whey was determined through Kjeldhal method as described in AOAC (2016). Ash contents of the fresh milk and cheese whey were checked by using method as described in AOAC (2016).

### 2.2. Preparation of ricotta cheese

Ricotta cheese was produced by the standard method of Wu *et al. (*2019) by using pasteurized milk in Dairy Laboratory. Ricotta cheese was prepared by heating raw milk to 63 ^0^C for 30 minutes and then cooling was done at 32 ^0^C. After cooling, 0.01% rennet was added to coagulate milk for the production of cheese whey. The cheese was then made by combining 80% cheese whey with 20% milk. It was acidified with citric acid until it reached a pH of 5.5. For coagulation, the acidified treatment was heated to 90°C for 30 minutes. Cooling was done at 20 ^0^C after coagulation and then drain the curd. The cheese was then kept at 4 degrees Celsius.

### 2.3. Edible coating preparation

Edible coating was prepared by following the approach of Kumar & Saini, 2021. In a 250 mL beaker, 5g whey powder was used as the basis material for the active edible coating for ricotta cheese, 2g xanthan gum was used to increase viscosity and kept at 40 °C for 90 min on magnetic stirrer. In a beaker, ten millilitres of glycerol were added as a plasticizer. Clove oil was added in different quantities in each solution for different treatments. The coating solution was homogenised by continuous stirring for 30 minutes using a magnetic stirrer. Different samples were prepared as shown in Table 1.

**Table 1:** Treatment Plan for coated ricotta cheese.

| Treatment (S) | Whey protein (g) | Clove oil % |
| --- | --- | --- |
| T<sub>0</sub> | - | - |
| T<sub>1</sub> | 2.5 | 0.1 |
| T<sub>2</sub> | 5.0 | 0.2 |
| T<sub>3</sub> | 7.5 | 0.3 |

T<sub>Rc</sub> = Ricotta cheese without any treatment
T<sub>Rc1</sub> = Ricotta cheese contains 0.1 percent clove oil and 2.5 gram of whey protein
T<sub>Rc2</sub> = Ricotta cheese contains 0.20 percent clove oil and with 5 grams of whey protein
T<sub>Rc3</sub> = Ricotta cheese contains 7.50-grams whey protein and 0.30 percent of clove oil

### 2.4. Edible coating analysis

#### 2.4.1. **pH**

pH of coating solution was determined by following the procedure as explained by Berti et al. (2019). Coating solution’s pH was determined using a digital pH metre. The pH metre was calibrated using buffers of pH 4 and 7. Following calibration, 20 ml of edible coating solution was placed in a beaker, and the electrode was dipped in the coating solution till consistent reading was obtained.

#### 2.4.2. Water activity

Water activity of edible coating solution was measured with water activity meter as shown by Berti et al. (2019). Water activity value was calculated by putting rod of water activity meter into beaker having 10 ml edible coating solution. Prior to measurement, the calibration was confirmed using saturated salt solutions. At each sampling location, the determination was made in duplicate during the ripening season.

### 2.5. Coating of ricotta cheese

Ricotta cheese was coated by following the method as explained by Ríos-de-Benito et al., 2021. Using a spray gun, the coating was applied to samples of Panela cheese; the samples were sprayed twice, separated by two minutes. All samples were drained on stainless steel screens after the coating process, air dried in a laminar flow cabinet for 20 minutes, and then placed into plastic “clam-shell” containers and stored at 4 ◦C. Physical, chemical, and microbiological tests on the cheese were performed after 0, 5, 10, and 15 days of storage. Cheeses that weren’t coated served as the control and were stored and examined simultaneously.

### 2.6. Physiochemical analysis

All physiochemical analysis of ricotta cheese was performed in Dairy Laboratory, Food Microbiology, and Biotechnology of NIFSAT, University of Agriculture, Faisalabad. Oven drying was used to determine moisture content Method No. 926.08 (AOAC, 2016). Whereas fat ×content was analyzed by using the butyrometer by using the procedure as described by the Siriwardana & Wijesekara, (2021). While protein content of ricotta cheese was determined using Kjeldalh’s method described by Margolies *et al*. (2018). The ash content of ricotta cheese was determined using Frau *et al*. (2014) method by using muffle furnace at 550°C until they reach a constant weight. Whilst Mileriene et al. (2021) method was used for the determination of pH of ricotta cheese. A portable pH meter was used for determining pH of samples.

### 2.7. Weight loss

Weight loss in cheese samples was determined by following the method as explained by Siriwardana & Wijesekara, (2021). Cheese samples were weighed individually on an automatic electric balance before and during storage. Relative weight loss was calculated in triplicate by using following formula:

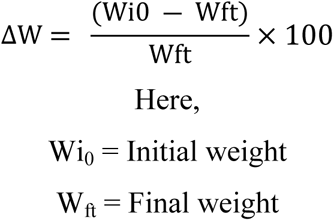

### 2.8. Anti-oxidant profile

#### 2.8.1. Diphenyl-2-picrylhydrazyl (DPPH) scavenging activity

The DPPH (1, 1-diphenyl-2-picrylhydrazyl) free radical scavenging activity of soft cheese extract was determined using the method as described by Masmoudi et al. (2020) with some modifications. Cheeses were examined for their DPPH scavenging activity immediately after preparing and during storage. By combining 3 g of cheese with a methanol:water (80:20; v/v) solution and stirring continuously for an hour at room temperature, phenolic extracts were produced. After the extract had been dried by evaporation at 40 °C the residues were suspended in methanol at a final concentration of 200 mg/ml. The mixture was vigorously shaken before being let to rest at room temperature in the dark. The absorbance was then determined at 515nm. The percentage reduction in DPPH due to a specific amount of each extract was computed and reported as a percentage reduction in radical scavenging percentage.

#### 2.8.2. TBARS

The amount of lipid oxidation in the treated cheese samples was determined using thiobarbituric acid reactive compounds (TBARS) by following the protocol as explained by Nourmohammadi et al. (2023). First of all, TBA (0.38%) and trichloroacetic acid (15%) were produced in 0.25 N HCl solution and added in a blender and homogenized at high speed for 3-4 minutes. A sample (5 ml) of the homogenate was incubated for color development for 15 minutes at 95°C in a water bath. The samples were centrifuged at 4500 g for 25 minutes after being refrigerated for 10 minutes. To measure the absorbance a spectrophotometer was used at a wavelength of 532 nm.

### 2.9. Microbial analysis

#### 2.9.1. Total viable count

Total viable count for each sample was determined by following the protocol explained by Mileriene et al. (2021). From each triplicate 10-g cheese sample was aseptically cut into three triangles to contain the cheese’s entire surface area and its center. Samples were put into a sterile stomacher bag, diluted in sterile Ringer’s solution (1:10, wt/vol), and homogenized for two minutes using a Stomacher 400 Circulator. The viable count agar was prepared and autoclaved for 15 minutes at 121°C. The numbers 10^-1^, 10^-2^, 10^-3^, 10^-4^, 10^-5^, 10^-6^, and 10^-7^ were written on seven sterilised test tubes. Each test tube received 9 mL of sterile Ringer’s solution. 1 g of homogenised cheese sample was transferred to the first test tube, and the contents were gently mixed together. The material was then transferred from the first to the second test tube and well mixed. Likewise, the sample from the second test tube was transferred to the third test tube. Other serial dilutions were made using the same process. 1 ml of each dilution’s contents was transferred to surface viable count agar plates, dispersed evenly, and incubated for 24 hours at 37 0C. With the use of a colony counter, the average number of colonies was counted from those dilutions that showed colonies ranging in size from 30 to 300.

### 2.10. Sensory evaluation

Sensory evaluation of coated cheese samples was performed according to the method as explained by Gulzar et al., (2020). A panel of five judges from NIFSAT, University of Agriculture Faisalabad, evaluated cheese samples for colour, taste, texture, and overall acceptability using a 9-point hedonic scale. Color, taste, texture, and overall acceptability were graded on a 9-point hedonic scale, with score 9 being great and score 1 being unsatisfactory. The total scores were calculated by putting all of the attribute scores together. Cheese samples were numbered and presented in a light to the panel members. After each sample, water was provided for a mouth rinsing.

### 2.11. Statistical Analysis

The findings of several studies were statistically evaluated at a 5 percent significance level according to Kontogianni et al. (2022). The statistical programmed statistical version 8.1 was utilized for investigation. The one-way analysis of variance was applied to find correlation between different parameters.

## 3. RESULTS AND DISCUSSION

### 3.1. Physicochemical analysis of fresh milk

The physico-chemical composition of the milk and whey samples are presented in Table 2. The results explicated that the protein, pH, acidity, fat, ash and moisture of raw milk and cheese whey were 3.14 & 2.62%, 6.52 &6.5, 0.17 & 0.11%, 4 & 0.21%, 0.61 & 0.53%, and 86.92 & 92.51%, respectively. The composition of milk was found slightly similar to the findings of Wu et al., (2019) who described the outcomes as pH (6.58%), acidity (0.15 %), fat (4.03%), protein (3.17%), moisture (87.94%), ash (0.66%). The cheese whey was obtained after curd formation during cheddar cheese production process. Its composition was found to be little similar to the results of (Wu et al., 2019) which are pH (6.49%), acidity (0.10%), fat (0.24%), protein (2.65%), moisture (92.55%), ash (0.59%).

**Table 2:** Chemical composition of milk.

| Components (%) | Milk | Whey |
| --- | --- | --- |
| Protein | 3.14 | 2.62 |
| Moisture | 86.92 | 92.51 |
| Acidity | 0.17 | 0.11 |
| Ash | 0.61 | 0.53 |
| Fat | 4 | 0.21 |
| pH | 6.52 | 6.5 |

### 3.2. Assessment of edible coating

The water activity, acidity, and pH of edible coating are presented in Table 3. The acidity and pH mean values were 0.12 and 6.70%, correspondingly, which were similar to results of Sudharani et al. (2021) which were 0.13% and 6.5 to 6.8 respectively. The average viscosity was found to be 3000 CP, which was in line with previous observations of Awad et al. (2005). A mean value of water activity (0.67%) was found which is closer to the findings of Beresford et al. (2001), who discovered that the water activities in edible plants were higher.

**Table 3:** Composition of edible coating.

| Components | Edible Coating |
| --- | --- |
| Viscosity (cP) | 3000 |
| Water Activity | 0.6 |
| Acidity (%) | 0.12 |
| pH | 6.7 |

### 3.3. Physicochemical Analysis of ricotta cheese

The mean values for the proximate content are shown in Table 4. Results showed that the moisture content in the ricotta cheese ranged from 63.250 to 73.37%. During storage study, T_Rc_ showed (73.360 to 63.250%), T_Rc1_ (73.360 to 64.237%), T_Rc2_ (73.370 to 66.243%), T_Rc3_ (73.360 to 68.217%). The results showed that group T_Rc3_ (0.30% clove oil and 7.50-g whey protein) had the best effects and less moisture loss during storage time. After storage, treatment T_Rc_ had lower moisture content than the other treatments. The overall findings for all groups showed a consistent decrease in moisture during storage. In comparison to the control sample, water loss was almost negligible in the clove oil and whey protein samples treated. The readings are in accordance with Hamdy *et al*. (2018), who also reported the decrease in moisture content in ricotta cheese. Humidity levels in Metata ayub soft cheese samples ranged from 46.70 to 50.30 percent after storage Eshetu & Asresie, (2019). Whereas the result showed that protein content of ricotta cheese was slightly increased. The protein content in ricotta cheese varies from 12.397 to 12.837. All treatments showed little increase in values of protein content such as T_Rc_ (12.397 to 12.82%), T_Rc1_ (12.72 to 12.817%), T_Rc2_ (12.73 to 12.837%), and T_Rc3_(12.727 to 12.83%) during 21 days of storage. This rise in protein concentration is mainly because of pH. Milk proteins coagulated best at pH 7, but as pH increased, protein denaturation occurred. Casein content, casein micelle characteristics (e.g., casein micelle size), and coagulation conditions (e.g., pH, temperature, and rennet concentration) have all been found to have an impact on the coagulation process. This could affect the way casein is arranged inside the protein matrix, as well as the final cheese’s microstructure and quality Lamichhane et al. (2018). The results are found in accordance with Hamdy *et al*. (2018) who reported the increase in protein level in ricotta cheese in 1-30 days of storage. Meanwhile the results regarding the fat content presented that mean value ranged from 8.12 to 8.23%. The highest fat contents were present in the T_Rc_ on the 21st day while the lowest fat content was observed in T_Rc1_ at 0^th^ day. T_Rc_ improved maximal fat content among the treated group, and an increasing trend of T_Rc1_>T_Rc3_>T_Rc2_ was seen and fat content decreased as storage time increased. This reduction in fat could be related to fat-protein interaction. Because T_Rc_3 has a larger protein content, more fat is associated to the protein, and its percentage drops. Vasiliauskaite et al. (2022) demonstrated the similar results and reported that fat content in cheese increased in cheese with storage time. The results are slightly similar to the readings of Hamdy *et al*. (2018) who reported the increase in fat level in ricotta cheese ranging from 15 to 15.15 during 1-21 days of storage.

**Table 4:** Physicochemical Analysis of ricotta cheese.

|  | **Days**<br>**(storage)** | **T<sub>Rc</sub>** | **T<sub>Rc1</sub>** | **T<sub>Rc2</sub>** | **T<sub>Rc3</sub>** |
| --- | --- | --- | --- | --- | --- |
| **Moisture** | 0 | 73.360 | 73.360 | 73.370 | 73.360 |
|  | 7 | 73.323 | 71.320 | 72.643 | 73.310 |
|  | 15 | 66.293 | 68.290 | 68.947 | 70.280 |
|  | 21 | 63.250 | 64.237 | 66.243 | 68.217 |
| **Crude**<br>**protein** | 0 | 12.397 | 12.720 | 12.730 | 12.727 |
|  | 7 | 12.750 | 12.757 | 12.760 | 12.750 |
|  | 15 | 12.780 | 12.770 | 12.783 | 12.790 |
|  | 21 | 12.820 | 12.817 | 12.837 | 12.830 |
| **Fat** | 0 | 8.1300 | 8.1200 | 8.1200 | 8.1200 |
|  | 7 | 8.1500 | 8.1300 | 8.1400 | 8.1333 |
|  | 15 | 8.1867 | 8.1600 | 8.1600 | 8.1700 |
|  | 21 | 8.2300 | 8.2000 | 8.2133 | 8.2200 |
| **Acidity** | 0 | 0.26 | 0.25 | 0.26 | 0.2567 |
|  | 7 | 0.2833 | 0.29 | 0.28 | 0.2733 |
|  | 15 | 0.31 | 0.2967 | 0.2967 | 0.3 |
|  | 21 | 0.32 | 0.31 | 0.32 | 0.3267 |
| **pH** | 0 | 6.45 | 6.52 | 6.57 | 6.63 |
|  | 7 | 6.37 | 6.21 | 6.29 | 6.16 |
|  | 15 | 5.91 | 6.01 | 6.11 | 6.40 |
|  | 21 | 5.51 | 5.63 | 5.82 | 6.36 |

The acidity in the ricotta cheese ranged from 0.25 to 0.3267%. When compared to T_Rc_ (Ricotta cheese without any treatment), the highest rising values were observed at 21 days of T_Rc3_ (Ricotta cheese contains 7.50-grams whey protein and 0.30 percent of clove oil), and the smallest growing tendency was seen at 0 days of T_Rc1_ (Ricotta cheese contains 0.1 percent clove oil and 2.5 gram of whey protein). T_Rc_3 has the largest influence on the acidity action of ricotta cheeses, followed by T_Rc_2 and T_Rc_1. The results are slightly similar to the readings of Hamdy *et al*. (2018) who reported the increase in fat level in ricotta cheese ranging from 15 to 15.15 during 1-21 days of storage. Dhuol and Hamid (2013) found that acidity increased from 0.41 to 1.14 percent at 0 day to 30 day storage, and finally 1.14+0.02 percent at 60 days of storage. The increase in cheese acidity has been reported by Ríos-de-Benito et al. (2021), which could be caused by temperature during storage. It also activated the raw milk natural micro-flora which generates acidity owing to fermenting lactose. That leads to a decrease in pH-value and lactic acid production. Whereas, the findings presented that as the storage period increases, the pH values of various treatments decrease. The most significant increases were identified after T_Rc3_ (0.3 percent clove essential oil and 7.5 gram whey protein) at 21 days, which was preceded by T_Rc2_(0.2 percent clove essential oil and 5 gram whey protein) treatment, while the least significant decreases were discovered after T_Rc_ treatment at 21 days. T_Rc3_>T_Rc2_>T_Rc_ was shown to be the general declining trend. The T_Rc3_ treatment, which includes 7.5 grams of whey protein and 0.3 percent clove oil as a coating, has a higher impact on soft cheese pH. The results are slightly similar to the readings of Hamdy *et al*. (2018) who reported the increase in pH level in ricotta cheese during 1-21 days of storage. The findings were in agreement with those of Mahmoudi *et al*. (2013), who discovered that utilization of Mentha longifolia during the ripening of probiotic feta cheese reduced the pH level from 4.53 to 4.44 after 60 days of storage at 4°C. He claimed that the lowering of lactic acid caused a considerable drop in pH during cheese storage.

### 3.4. Analysis of water activity and weight loss

The mean values of the water activity were presented in Table 6. The result explicated that the water activity ranged from 0.90 to 0.97 in the ricotta cheese. The maximum value (0.96) of the water activity was observed in the **T_Rc_**on the 0th day whereas, minimum water activity (0.90) was explored in the **T_Rc_** on the 21st day. Water activity reduced as storage duration increased. At 21 days, T_Rc_ has the lowest value, followed by T_Rc2_, while T_Rc3_ had the lowest reducing values at 21 days. Water activity is an important factor in determining the life span of food items, such as higher the water activity, the higher the microbial deterioration. Water activity is also determined by the moisture level of the product. The findings are in agreement with those of Pluta-Kubica et al. (2020) who observed that there was a reduction in water activity in all cheese samples throughout storage. The mean values of weight loss ranges from ricotta cheese groups. The results reveal that there are substantial disparities between the various treatment groups. The results of the research reveal that there are irregular variations in soft chess weight loss between treatments. With increased storage duration, weight loss decreases. The highest reduction was seen after 21 days of therapy, while the smallest reduction was seen after 0 days of treatment. T_Rc_3>T_Rc_2>T_Rc_1 was detected as 8u9 y67a decrease Percent within the same day. Consequently, weight loss decreases as storage duration increases, with the greatest drop occurring after 21 days of T_Rc_3. During storage, the overall values for all treatments exhibited continuous weight loss. Clove essential oil 0.3% and 7.5 g whey protein treated sample resulted in little weight reduction. Whey protein reduces the water loss in cheese due to its gas barrier properties. Evaporation of water vapors is done during the storage of cheese, which is affected by packing material. The results of this experiment are consistent with those of Siriwardana & Wijesekara, (2021) who found that soft cheese loses 46.4-50.2 pounds during cold storage. During the storage of soft cheese samples, weight loss was also observed (46.7-50.3 percent) Eshetu & Asresie, (2019).

**Table 5:** Analysis of water activity and weight loss.

|  | Days<br>(storage) | T<sub>Rc</sub> | T<sub>Rc1</sub> | T<sub>Rc2</sub> | T<sub>Rc3</sub> |
| --- | --- | --- | --- | --- | --- |
| **Water activity** | 0 | 0.97 | 0.96 | 0.95 | 0.95 |
|  | 7 | 0.94 | 0.94 | 0.95 | 0.94 |
|  | 15 | 0.95 | 0.93 | 0.92 | 0.92 |
|  | 21 | 0.90 | 0.91 | 0.91 | 0.92 |
| **Weight loss** | 0 | 0.031 | 0.021 | 0.021 | 0.021 |
|  | 7 | 3.5 | 3.09 | 2.87 | 2.11 |
|  | 15 | 19.35 | 13.91 | 12.63 | 11.87 |
|  | 21 | 26.6 | 24.69 | 23.57 | 22.64 |

**Table 6:**
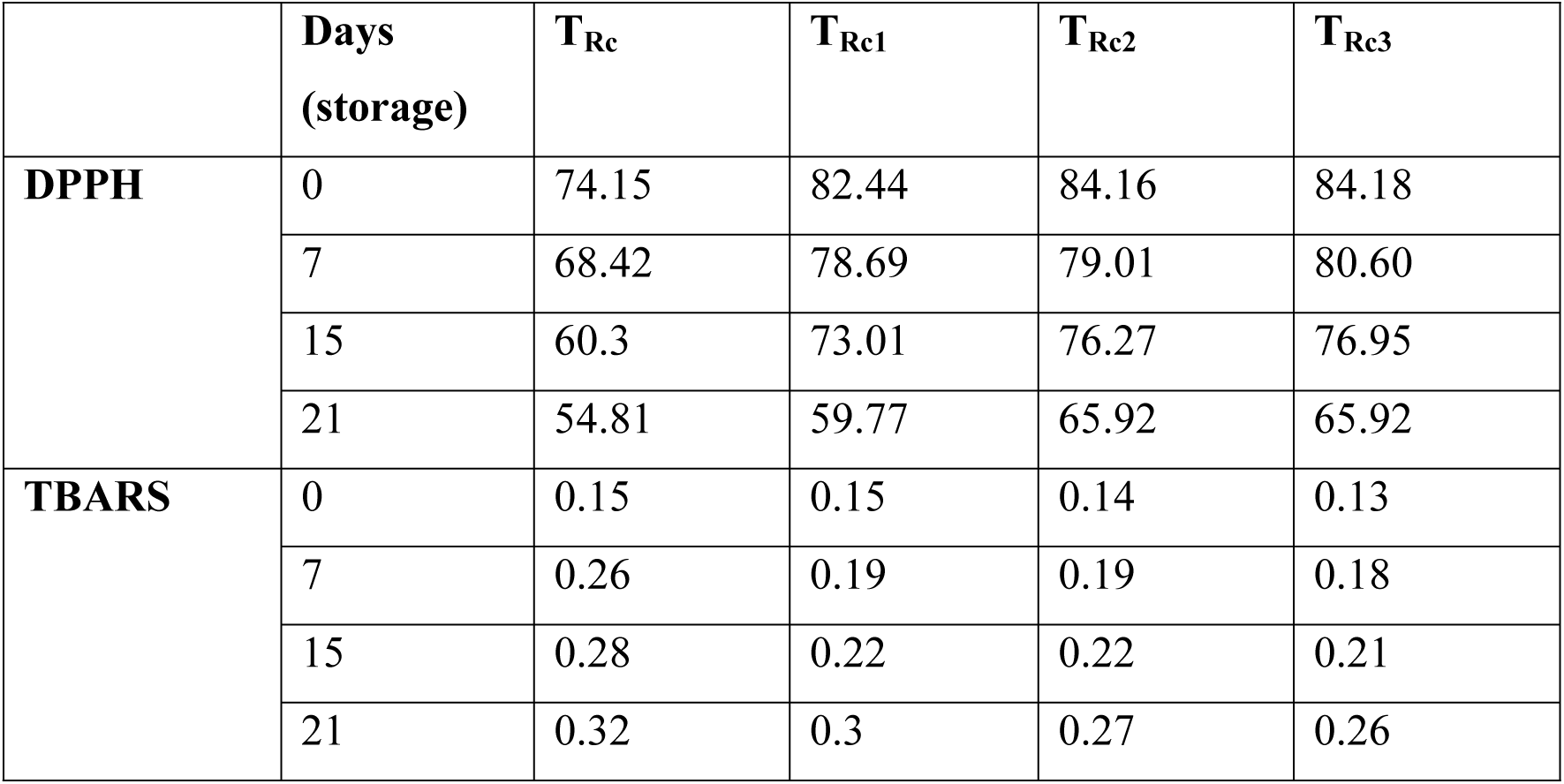
Antioxidant profile of ricotta cheese.

|  | **Days**<br>**(storage)** | **$T_{Rc}$** | **$T_{Rc1}$** | **$T_{Rc2}$** | **$T_{Rc3}$** |
| --- | --- | --- | --- | --- | --- |
| **DPPH** | 0 | 74.15 | 82.44 | 84.16 | 84.18 |
|  | 7 | 68.42 | 78.69 | 79.01 | 80.60 |
|  | 15 | 60.3 | 73.01 | 76.27 | 76.95 |
|  | 21 | 54.81 | 59.77 | 65.92 | 65.92 |
| **TBARS** | 0 | 0.15 | 0.15 | 0.14 | 0.13 |
|  | 7 | 0.26 | 0.19 | 0.19 | 0.18 |
|  | 15 | 0.28 | 0.22 | 0.22 | 0.21 |
|  | 21 | 0.32 | 0.3 | 0.27 | 0.26 |

### 3.5. Antioxidant Activity

The mean values for the antioxidant contents are shown in Table 5. Results showed that the DPPH content in the ricotta cheese ranged from 54.81 to 84.18. During the storage period, the T_Rc3_ (7.5 g whey protein and 0.3% clove oil) has the highest DPPH values in ricotta cheese at 0 days while the T_Rc_1 has the lowest DPPH levels after 21 days. The findings show that as the storage period increases, antioxidant capacity decreases. However, an uneven pattern was found when different treatments of ricotta cheese were stored for varying amounts of time. The declining pattern was noticed as T_Rc3_>T_Rc2_>T_Rc1._ DPPH method is used to determine antioxidant activity based on the process by which antioxidants prevent oxidation of lipid, resulting in DPPH radical scavenging. It also predicts the free radical scavenging capacity. It is also preferable because it is completed in a very short duration. They are also utilized to keep food fresh for longer. Antioxidants included in ricotta cheese are utilized to keep food fresh for longer. The results are slightly similar to the readings of Hamdy *et al*. (2018) who reported the decrease in antioxidant level in ricotta cheese during 1-21 days of storage.

The findings regarding Thiobarbituric acid reactive substances (TBARS) reveal that the mean values of the various treatment groups varied significantly. The results demonstrate that when the storage duration was extended, the Thiobarbituric acid reactive substances values increase. The greatest values were found at day 21 of T_Rc_, followed by T_Rc1_, and the smallest values were found at day 21 of T_Rc3_. T_Rc_>T_Rc1_>T_Rc2_ > T_Rc3_ was shown to be the overall trend of decline in values. TBARS assay is commonly used to measure lipid oxidation in soft cheese under storage conditions. The TBARS value in soft cheese is a sign of oxidative degradation. Esparvarini et al. (2022) also investigated the rise in Thiobarbituric acid reactive substances reactive material values at refrigeration temperature during storage period. The results are slightly similar to the readings of Hamdy *et al*. (2018) who reported the increase in Thiobarbituric acid reactive substances level in ricotta cheese during 1-21 days of storage.

### 3.7. Microbiological analysis

#### 3.7.1. Total Viable count

The total viable count (cfu/g) is microbiological technique which is used to estimate the microbial concentration such as yeast, mold and bacteria. The total viable count declined as lactic acid bacterial production increased, possibly due to synergistic effect. Increased production of lactic acid will reduce dangerous microorganisms and raise the level of acidity of the product, possibly resulting in product preservation. With the passage of time, the total viable count may continue to decline. However, Mileriene et al. (2021) reported that after a certain number of days, yeast production may cause the product to deteriorate but to a lesser extent because coating will prevent the yeast growth.

The results are significant (P0.05) according to the assessment. The findings determine that the means of various groups differed substantially. Table 7 shows the impact of various treatments and storage durations on the total viable count of ricotta cheese. T_Rc_3 (7.50-grams whey protein and 0.30 percent of clove oil) in ricotta cheese showed the minimum rise in total viable count due to strong antimicrobial effect of clove oil during 0-21 days of storage. T_Rc_3 showed the best results than other treatments (T_Rc2_, T_Rc1,_ T_Rc_) respectively. Overall, the findings show that treatment decreasing pattern of the total viable count was T_Rc3_>T_Rc2_> T_Rc1_ >T_Rc._

**Table 7:** Microbiological analysis of ricotta cheese.

| **Treatment** | **0-day** | **7-days** | **15-days** | **21-days** |
| --- | --- | --- | --- | --- |
| **$T_{Rc}$** | 5.6033 | 5.8100 | 5.9433 | 6.1067 |
| **$T_{Rc1}$** | 5.4867 | 5.5933 | 5.7733 | 5.8100 |
| **$T_{Rc2}$** | 5.4467 | 5.5933 | 5.6300 | 5.7200 |
| **$T_{Rc3}$** | 5.4167 | 5.4900 | 5.5333 | 5.5800 |

The findings were found similar with those of Siriwardana & Wijesekara, (2021) who observed that the total viable count of cheese dropped during cold storage from the production day.

### 3.8. Sensory Analysis

According to the research, the results for texture analysis are significant (P<0.05). The results demonstrate that the mean values of different treatments varied significantly. The results reveal that as storage time increased, the texture values of various treatments dropped. The findings are in agreement with those of Jalilzadeh et al. (2020) who observed that soft cheese texture degaraded throughout storage.

Results for taste are significant (P<0.05), according to the analysis. There are substantial variations in the means of the various groups. The results show that when the storage duration was increased, the values fell. The findings are in agreement with those of Nourmohammadi et al. (2023) who observed that soft cheese taste loses throughout storage.

The findings for color analysis are significant (P<0.05) according to the analysis. The means data reveal that the means of the several groups varied significantly. The interplay of treatments and storage on the color of ricotta cheeses is depicted in the table. The results show that the values dropped as the storage period increased, with the highest values at T_Rc3_ and the lowest values at T_Rc1_. The patterns at T_Rc3_ 0 day>7 days>15 days>21 days showed that T_Rc3_ values declined with a greater storage period. T_Rc3_ has better color than T_Rc1_ and T_Rc2,_ according to the results obtained. The findings are in agreement with those of Jalilzadeh et al. (2020) who observed that soft cheese colour loses throughout storage.

The table shows the impact of treatments and storage time on overall acceptance. Overall, the effects of storage and treatment on ricotta cheese acceptability were significant (P<0.05). During the storage periods, T_Rc_3 (whey protein 0.75 percent, clove oil 0.3 percent) from the control group had the highest acceptability range and T_Rc_1 (whey protein 2.5g, clove oil 0.1 percent) had the lowest acceptability area. The findings also show that when the storage duration was increased, the acceptability dropped. Because of the various components of different therapy groups, there is a disparity in acceptance. The results of this experiment are consistent with those of Arshad *et al,* (2020), who found that ricotta cheese had an overall acceptance of 8.4-5.10 at refrigerated temperature.

**Table 8:** Sensorial characteristic of ricotta cheese.

| **Sensory Analysis** | **Days (Storage)** | **T<sub>Rc</sub>** | **T<sub>Rc1</sub>** | **T<sub>Rc2</sub>** | **T<sub>Rc3</sub>** |
| --- | --- | --- | --- | --- | --- |
| **Texture** | 0 | 6.93 | 6.79 | 6.53 | 7.64a |
|  | 7 | 6.21 | 6.43 | 6.39 | 7.32 |
|  | 15 | 6.11 | 6.200 | 6.151 | 7.22c |
|  | 21 | 6.05 | 6.131 | 6.101 | 7.10 |
| **Taste** | 0 | 6.43 | 6.65 | 6.75 | 7.61 |
|  | 7 | 6.22 | 6.32 | 6.42 | 7.30 |
|  | 15 | 5.50 | 6.26 | 6.19 | 7.11 |
|  | 21 | 5.41 | 6.24 | 6.25 | 7.02 |
| **Color** | 0 | 6.67 | 6.78 | 7.24 | 7.51 |
|  | 7 | 6.341 | 6.31 | 7.11 | 7.31 |
|  | 15 | 6.10 | 6.20 | 7.04 | 7.13 |
|  | 21 | 6.05 | 6.15 | 6.98 | 7.04 |
| **Overall acceptability** | 0 | 7.661 | 7.96 | 8.02 | 8.45 |
|  | 7 | 7.52 | 7.42 | 8.21 | 8.25 |
|  | 15 | 7.20 | 7.21 | 8.01 | 7.85 |
|  | 21 | 7.14 | 7.13 | 7.98 | 7.80 |

## Conclusion

Different concentrations of whey protein and clove oil were employed in coating to observe their effects on ricotta cheese. Storage conditions had a significant (p<0.05) effect on physiochemical properties of ricotta cheese such as protein, ash, moisture, fat, pH and acidity. Ricotta cheese showed a significant (p<0.05) decrease in the total viable count (TVC). During storage, there is a significant decrease in the values of DPPH radical scavenging activity in coated ricotta cheese from 1-24 days of storage. However, the radical scavenging activity increased as clove essential oil concentration increased. Sensory evaluation of ricotta cheese reported that the storage effect on all sensory parameters was significant (p<0.05). Clove oil treated samples of ricotta cheese showed different results. Hence, clove essential oils and whey protein maximum concentrations in coating leads to high quality ricotta cheese.

## Credit Authorship Contribution Statement

Muhammad Aamir, Muhammad Afzaal designed the study and Eram Sarwar had conducted under the supervision of Muhammad Aamir and Muhammad Afzaal. Ali Ikram, Huda Ateeq and Faiza Kiran performed the study and participated in drafting the article with Noor Akram. Farhan Saeed and Aftab Ahmad helped in developing the whole concept and editing. Noor Akram, Ali Ikram and Huda Ateeq helped in preparing Figures and Tables, the overall quality of the manuscript was maintained by Aftab Ahmed. Farhan Saeed and Muhammad Afzaal wrote, edited and revised the manuscript critically. Shahzad Hussain and Mahbubur Rehman Khan revised the final written paper. The final version of the manuscript has been read and approved by all listed authors.

## Data Availability

Even though adequate data has been given in the form of tables and figures, however, all authors declare that if more data is required then the data will be provided on a request basis.

## Acknowledgment

The authors appreciate the support from the Researchers Supporting Project number (RSPD2023R1073), King Saud University, Riyadh, Saudi Arabia.

